# CD200 depletion in glioma enhances antitumor immunity and induces tumor rejection

**DOI:** 10.1101/2024.09.08.611922

**Authors:** Itay Raphael, Anzar A. Mujeeb, Elisabet Ampudia-Mesias, ReidAnn E. Sever, Brandon McClellan, Stephen C. Frederico, Chaim T. Sneiderman, Apoorva Mirji, Ali Daba, Francisco Puerta-Martinez, Michal Nisnboym, Wilson B. Edwards, Michael Graner, Christopher L. Moertel, Maria G. Castro, Gary Kohanbash, Michael R. Olin

**Author notes:** Co-corresponding authors: <u>Gary Kohanbash</u>: University of Pittsburgh School of Medicine, UPMC Children’s Hospital of Pittsburgh, 7128 Rangos Research Building, 530 45th St., Pittsburgh, PA 15201, USA.; <u>Michael R. Olin</u>: University of Minnesota, 2-167 Moos Tower, 515 Delaware St SE, Minneapolis, MN 55455, USA. Co-first authors (Itay Raphael, Anzar A. Mujeeb). **Competing interests**: Drs. Olin and Moertel hold equity in OX2 Therapeutics, which holds licensing rights to the development of CD200 immune checkpoint inhibitor. These interests have been reviewed and managed by the University of Minnesota in accordance with its Conflict-of-Interest policies.

## Abstract

High-grade gliomas are a major health challenge with poor prognosis and high morbidity. Immune-checkpoint inhibitors (ICI) have emerged as promising therapeutic options for several malignancies yet show little efficacy against central nervous system (CNS) tumors. CD200 is a newly recognized immune checkpoint that modulates immune homeostasis. CD200 protein is expressed by a variety of cells, including immune cells and stromal cells, and is overexpressed by many tumors. The shedding of CD200 from tumor cells can create an immunosuppressive environment that dampens anti-tumor immunity by modulating cytolytic activity and cytokine expression both within and outside the tumor microenvironment (TME). While it is well-accepted that CD200 induces a pro-tumorigenic environment through its ability to suppress the immune response, we sought to determine the role of glioma-specific expression of CD200. We show that CD200 is expressed across glioma types, is shed from tumor cells, and increases over time in the serum of patients undergoing immunotherapy. Using CD200 knockout (KO) glioma models, we demonstrated that glioma cell-derived CD200 promotes tumor growth in vivo and in vitro. Notably, CD200 KO gliomas are spontaneously rejected by their host, a process that required a fully functional immune system, including NK and T-cells. Moreover, we report that glioma-derived or brain-injected soluble CD200 contributes to the suppression of antigen-specific CD8 T-cells in the draining lymph nodes (dLNs). Our work provides new mechanistic insights regarding CD200-mediated immunosuppression by gliomas.

**Statement of significance:** We demonstrate mechanisms of the druggable glioma-derived CD200 checkpoint on tumor growth and immune suppression.

## Introduction

Glioblastoma (GBM; IDH-WT) is the most malignant primary brain tumor in adults, with median overall survival of 12-18 months and the 5-year survival rate is around 7% (1).

Immunotherapy has emerged as an important therapeutic option that can supplement standard of care (surgical resection and chemo-radiotherapy) for achieving long-term survival. However, developing successful immunotherapy approaches for gliomas has been hampered by an immunosuppressive tumor microenvironment (TME). Within the complex mechanisms of TME immunosuppression, immune-checkpoint molecules, typically responsible for maintaining self-tolerance and regulating immune responses, are exploited by tumors to evade immune destruction (2,3). Thus, immune-checkpoint inhibitors (ICIs) have emerged as a potential therapeutic option to block the inhibitory signals and to promote anti-tumor immunity (4). To-date, ICIs demonstrated survival benefits in specific solid tumors with limited impact on patients with GBM (5). ICIs have had a broad impact; several antibodies targeting cytotoxic T lymphocyte antigen-4 (CTLA4) or program death-1 (PD-1) and its ligand PD-L1 have been approved for use in select cancers (6). Additionally, tumors may utilize more than one immune-checkpoint to their benefit, severely impeding the efficacy of monotherapy with ICIs and frequently requiring the use of multiple ICIs to achieve efficacy (7).

A key player in immune modulation is the glycoprotein CD200, also known as OX-2 membrane glycoprotein, now recognized as an established immune-checkpoint molecule (8). CD200 is primarily expressed throughout the CNS but is also reported to be expressed on immune cells and some endothelial cells (9,10). Additionally, CD200 is upregulated on certain tumors and on tumor vascular endothelial cells (11). CD200 serves a pivotal role in maintaining immune tolerance and protects healthy tissues from immune-mediated damage through binding with its immune inhibitory receptor (CD200R1) (12). Similar to other immune checkpoint proteins, CD200 plays a pro-tumorigenic role in many tumor types, primarily by suppressing anti-tumor T-cell and natural-killer (NK) cell responses (13–16). Additionally, evidence from prior studies demonstrated that targeting CD200 enhances immunotherapy outcomes (17–19). Furthermore, a clinical trial evaluating an anti-CD200 antibody, samalizumab, demonstrated a conferred therapeutic benefit to a subset of patients and highlighted the potential of targeted CD200 inhibition (19). A plausible explanation for the observed shortcomings of samalizumab in non-responding patients is that the multiple mechanisms of the pro-tumorigenic role of CD200 extend beyond direct suppression of the anti-tumor T-cell response not susceptible to the CD200 inhibition (20).

Despite the evidence that CD200 plays a significant role in the immune suppressive environment and promotion of tumor growth, the role of tumor-derived CD200 in gliomas remains unresolved. Here, we show that CD200 is expressed across glioma types, is shed from glioma cells, and increases over time in patients’ serum. Furthermore, we developed a mouse syngeneic CD200 KO glioma model and used it to demonstrate that glioma-derived expression of CD200 enhances glioma cell growth and is a major contributor to the immunosuppression of NK cells and antigen-specific CD8 T-cells. Our data reveal that CD200 KO gliomas were spontaneously rejected by their host, and this rejection process depends on a functional immune system.

## Materials and Methods

### Brain tumor inoculation and in vivo experiments and imaging

Mice were anesthetized using ketamine (75mgkg) and dexmedetomidine (0.5mg/kg) before stereotactic implantation with GL261 WT and GL261 CD200 KO cells in the right striatum. The coordinates for implantation were 1.0 mm anterior and 2.0 mm lateral from the bregma, and 3.5 mm ventral from the dura. Tumor neurospheres were injected at a rate of 1 μL/min. Mice were given a combination of buprenorphine (0.1mg/kg,S.C) and carprofen (5mg/kg, subcutaneously) for analgesia. A single dose of 14×10³ wildtype GL261-Luc+ cells and GL261-CD200 KO-Luc+ cells was used for tumor inoculation. Tumor burden was determined by bioluminescent imaging, using 150 mg/kg d-luciferin potassium salt (BioVision), and quantifying light emitted from tumor tissue using a Xenogen IVIS System (Perkin Elmer, Inc., MA). Images were acquired using auto exposure settings (exposure time 1-200 sec, binning 4/8/16, field of view 23 cm, f/stop 1, emission filter open) and signal was measured and recorded as total flux (photons/sec). Collection of tissue and cells (e.g., spleens and lymph-nodes) was performed following perfusing with Tyrode’s solution. In selected experiments, C57BL/6J mice were implanted with ALZET pumps for a seven-day infusion of CD200 protein. The pump was placed at the same location as glioma cells for delivery. Dexamethasone Treatment of Mice: 6–8-week-old male C57BL/6 mice were injected intraperitoneally with 2.5 mg/kg/day pharmaceutical-grade dexamethasone (R&D Systems) or vehicle (saline) for 3 consecutive days prior to stereotactic implantation of 45,000 bioluminescent CD200KO GL261 cells on day 4. Mice were imaged weekly after day 8. The mice were housed in the Animal Facility at the University of Minnesota. All experimental procedures were conducted in adherence to national guidelines and regulations, with prior approval from the University of Minnesota Institutional Animal Care and Use Committee (IACUC).

### Cell lines and culture conditions

Murine cancer cell line GL261 stably expressing luciferase, was purchased from the American Type Culture Collection (Manassas, VA). CD200 knockout GL261 cells (CD200KO GL261) were in-house generated using the CRISPR/Cas9 system. Cells were cultured in Dulbecco’s modified Eagle’s medium-DMEM supplemented with 10% FBS (Gibco, ThermoFisher Scientific) and 100 μg/mL Penicillin/Streptomycin (Corning, Inc., NY) at 37°C, 5% CO2 and 95% humidity. All cell lines were mycoplasma-free.

### Generation of a GL261-CD200KO cell line

CD200 Knockout GL261 cells were generated using the CRISPR/Cas9 system. Cas9/EGFP plasmid PX458 was purchased from ADGENE (#48138) and used for transient expression of Cas9. The guide sequences targeting CD200 genomic loci were designed using the online bioinformatics tool ’CRISPOR’. Neon® electroporation-mediated transfection system (ThermoFisher Scientific) was utilized for transfection of Cas9/GFP plasmid and gRNA into GL261 cells. Briefly, GL261 cells were grown in Dulbecco’s modified Eagle’s medium-DMEM supplemented with 10% fetal calf serum, 1% penicillin/streptomycin, and incubated at 37°C until 80% confluent. Cells were harvested by trypsinization, washed in DMEM, and resuspended in 1X PBS at a concentration of 5e10^6^ cell density (cells/ml). 100 μl of the cell suspension was used for the transfection. Cells were then precipitated and resuspended in 10 μl Neon Buffer R, mixed with 0.5 μg of each CRISPR/Cas9 vector and 1 μl (100 pmol/μl) sgRNA Synthego (Synthego) and incubated for 2 min. Electroporation was conducted using 1,425 volts, 30 milliseconds, and 2 pulses. Post-transfection, cells were placed in a 6-well plate with DMEM lacking antibiotics and cultured for 48-72 hours. Viable cells were then sorted three times using a FACSAria II cell sorter (BD Biosciences). PCR and Western blot assays confirmed gene removal, with non-transfected and empty plasmid-transfected cells employed as controls.

### Western blot analysis

Wildtype GL261 or CD200KO GL261 cells were grown in a 6 well plate with 3 mL of supplemented DMEM until 80% confluence was observed. Cells were then lysed on ice for 10 minutes in 200 µl of RIPA buffer (ThermoFisher Scientific). Protein concentrations were determined using the bicinchoninic acid colorimetric method (Pierce Biotechnology, Waltham, MA). Lysates were diluted in reducing sample buffer (Novex), and 50 µg were loaded per lane on a 4 to 12% SDS-PAGE gel (Nu-Page) and run at 160 V (0.8 V h). Gels were then transferred to PVDF membranes (BioRad, Hercules, CA) at 25V using a semi-dry transfer system (Invitrogen), blocked using 5% nonfat dry milk/0.05 mM Tris-buffered saline with 0.05% Tween 20 for 1 h, incubated overnight at 4 °C with an anti-OX2 antibody, 200 μg/mL (1:1000, Santa Cruz Biotechnology, Dallas, TX) in blocking buffer, washed 3 times in TBS/Tween 20 over 15 min, and incubated with a secondary HRP-conjugated antibody for 1 h at room temperature (1:10,000, Jackson ImmunoResearch). Protein bands were visualized by ECL Plus Chemiluminescent Substrate (Pierce Biotechnology) using an iBright Imaging Systems (ThermoFisher Scientific).

### Cell proliferation assays

2.5×10^5^ wildtype or CD200KO GL261 cells were plated and monitored for cell growth. Cells were then stained with 0.4% trypan blue (Invitrogen) and counted using a Neubauer cell counting chamber. All experiments were performed at least twice with two-three replicates per experiment.

### Natural Killer (NK) cells depletion

Mice were injected intraperitoneally (IP) with 100μg monoclonal anti-NK1.1 (Purified anti-mouse NK-1.1 Antibody BioLegend 108702) diluted to a final volume of 400μL in (DPBS (Cat# 14190250, Gibco), pH 7.4) two days before the isolation of the spleen for the analysis of splenocytes. Mice were euthanized with isoflurane, and the spleens were collected. The spleen was homogenized using a syringe plunger and passed through a 70um strainer to collect the splenocytes. The strainer and plunger were rinsed with 5-8 mL of media (RPMI-1640 (Cat#11875119, Gibco). These steps were repeated to ensure maximum splenocyte collection. The tube containing the splenocytes was centrifuged at 1500 rpm for 5 min at 4 C. The supernatant was discarded in bleach, and the pellet was resuspended in 1X RBC lysis buffer for 60-90 seconds. The RBC Lysis buffer was neutralized with RPMI, 10 times the volume of the lysis buffer. The tubes were centrifuged one more time using the same settings as before. The supernatant was removed, and the white pellet consisting of splenocytes was obtained for further analysis.

### Flow Cytometry Analysis

Spleens were collected and homogenized using Tenbroeck (Corning) homogenizer in DMEM (cat# 12430054, Gibco) media containing 10% FBS (cat#10437028, Gibco). Cells were resuspended in PBS containing 2% FBS, and non-specific antibody binding was blocked with FC block. All stains were carried out for 30min at 4°C and washed with flow buffer between each staining step. NK cells were gated as CD45^+^, CD3^-^, and NK1.1^+^. Data acquisition was performed using FACS ARIA II (BD Biosciences) and analyzed using FlowJo software.

### RNAseq analysis and Gene Set Enchainment Analysis (GSEA)

RNAseq reads (from fastq files) were aligned with the mouse genome (Mus musculus genome assembly) using Salmon (v0.9.1). Transcript-level quantification was collapsed onto gene-level quantification using the tximport package in R based on the gene definitions provided by the same ensemble release (Ensembl transcript annotations. We then performed Deseq2 to identify differentially regulated genes and normalized gene counts (transcript per million; TPM). GSEA was performed on pre-ranked gene list based on Deseq2 data. Each gene was assigned a rank based on the FDR-adjusted *p* values (adj *p*) and correlation coefficients (*r*) or fold-change (FC). A final gene rank list from each analysis was used to perform pre-ranked Gene Set Enrichment Analysis (GSEA) (https://www.gsea-msigdb.org/) using C5 Biological Process ontology gene sets of the Molecular Signatures Database (MSigDB) v3.0. GSEA network analysis was performed using the EnrichmentMap plugin (21) on Cytoscape platform (22) with adjustments for *p* value cutoff < 0.001, FDR Q-value cutoff < 0.1, and gene-set overlap coefficient of 0.4. Node colors were set to represent the gene-set normalized enrichment scores; where red nodes are positive scores (positively enriched in CD200 KO tumors), and blue nodes are negative scores (negatively enriched in CD200 KO tumors). Node sizes were set to denote the gene-set size. TCGA-generated glioblastoma (GMB) and low-grade glioma (LGG) tumor bulk RNAseq data (23) analysis and visualization were performed using GlioViz (http://gliovis.bioinfo.cnio.es/) (24) and GEPIA2 (http://gepia2.cancer-pku.cn/) (25) platforms using default settings.

### Statistical analysis

Sample sizes were selected based on data from previous experiments done in our laboratories and published results from the literature. Animal experiments were performed after randomization. Kaplan-Meier survival curves were assessed using the log-rank (Mantel-Cox) test with Prism 8.1 (GraphPad Software). P values less than 0.05 were considered statistically significant.

## Results

### CD200 expression differs between glioma subtypes and correlates with prognosis

The inhibitory CD200 protein is highly expressed in a variety of human tumors (26,27) and was reported to be shed by tumor cells and to suppress antigen presentation and T-cell responses (27). Analysis of CD200 RNA expression within gliomas demonstrated that CD200 is expressed across all major histology groups at various levels (**Fig. 1a**). We noted significant differences in the mean expression levels of CD200 between histology groups, in which CD200 expression trended to be expressed at lower levels in GBM (**Fig. 1b**). Stratification of patients into groups based on the expression of CD200 demonstrated that low CD200 expression was associated with poor prognosis of patients, including worse overall survival rate and poor disease-free survival rate (**Fig. 1c-d**). Together, these data suggest that CD200 may be involved in the regulation of glioma immunity merit further investigation.

**Figure 1:**
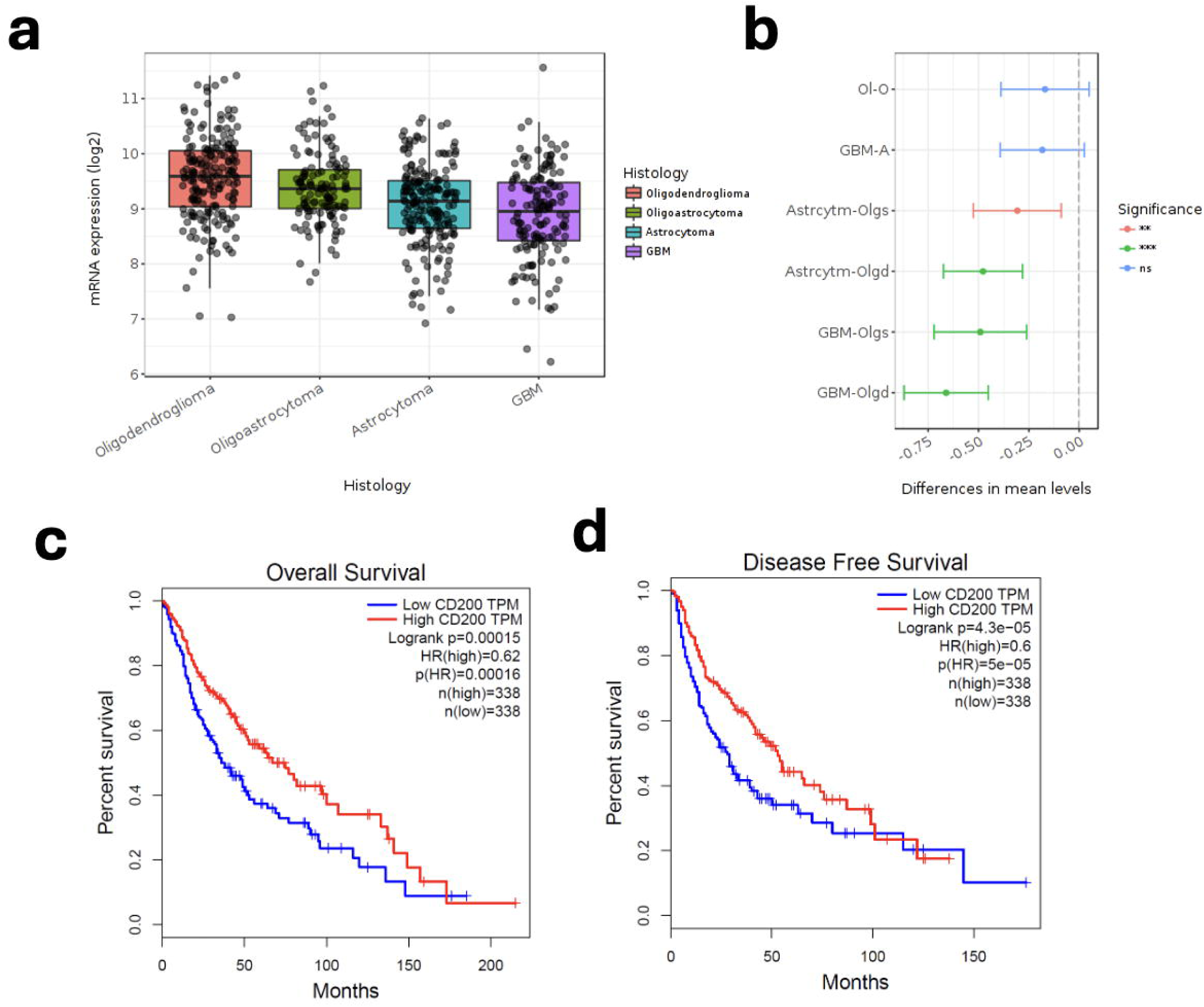
CD200 gene expression is associated with histological grade and patient prognosis in glioma. **a**. CD200 RNA expression across glioma histological types. **b**. Pairwise comparisons in CD200 RNA expression between glioma histological types. Ol-o: Oligoastrocytoma vs. Oligodendroglioma; GMB-A: GBM vs. Astrocytoma; Astrocytm-Olgs: Astrocytoma vs. Oligoastrocytoma; Astrocytm-Olgd: Astrocytoma vs. Oligodendroglioma; GBM-Olgs: GBM vs. Oligoastrocytoma; GBM-Olgd: GBM vs. Oligodendroglioma. P values were determined using Tukey’s Honest Significant Difference (HSD) test **c-d**. Kaplan–Meier survival curves of glioma patients showing overall survival (**c**) and progression free survival (**d**) stratified by CD200 expression (high=red vs. low=blue). P values and hazard ratio were determined using a log-rank test.

### Reducing CD200 expression in glioma cells improves mouse survival and is associated with changes in cellular function and immune modulation gene networks

To study the role of CD200 in glioma, we knocked out (KO) the *Cd200* gene in GL261 murine glioma cells, a well-established preclinical model of GBM. CD200 protein expression was assessed using Western blot and flow cytometry. GL261 CD200-KO cells presented with no observed CD200 expression as compared with wildtype (WT) cells (**Fig. 2a, b**). We next investigated the role of CD200 expression in glioma cells in vivo. Strikingly, mice bearing CD200 KO gliomas spontaneously rejected CD200 KO tumor cells and presented with significantly improved survival rates as compared to mice inoculated with WT gliomas (**Fig. 2c**). Transcriptomic analysis of these tumors demonstrated significant changes in gene expression profiles (**Fig. 2d**). Pathways analysis of the differently regulated genes demonstrated that CD200 KO tumors upregulated genes associated with immune cell responses, cellular organization, tissue development, and chemotaxis (**Fig. 3e**). In contrast, downregulated genes in CD200 KO tumors were associated with cellular respiration and metabolism (**Fig. 3e**). Among top regulated pathways we noted cytokine production, extracellular matrix assembly, angiogenesis, and chemotaxis -related pathways positively enriched in CD200 KO tumors, while oxidative phosphorylation, mitochondrial functions, and energy/ATP metabolism -related pathways were enriched in the CD200 WT tumors (**Fig. 2f**). Gene ontology analysis of genes positively correlated (*r* >0.3, adj p<0.05) with CD200 in GBM demonstrated that CD200 expression was associated with several neuronal development and signaling pathways including neurotransmitter and ion channel activity, neuronal development, and tissue morphology (**Fig. 2g**). Together these data demonstrate that CD200 has an intrinsic role in regulating glioma development and its cellular function, including cell cycle and immunity.

**Figure 2:**
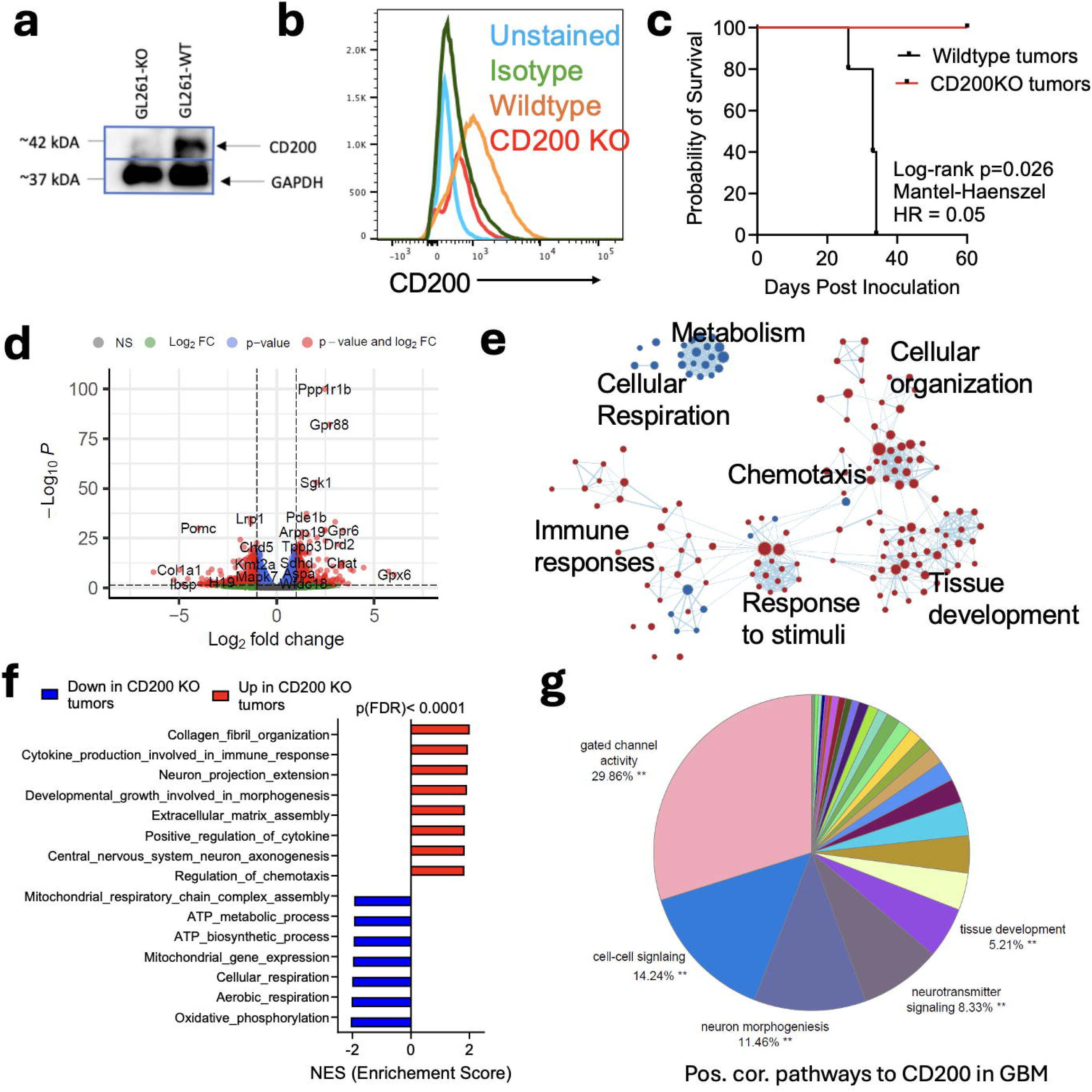
Differential gene expression and pathway analysis of CD200 knockout (KO) gliomas. **a-b**. CD200 protein expression in wildtype (WT) and CD200 KO GL261 glioma cells determined by (**a**) Western-blot and (**b**) flow-cytometry analysis. **c**. Kaplan–Meier survival curves curve of mice inoculated with WT or CD200 KO gliomas. p values were determined using a log-rank test. Hazard ratio was determined using Mantel-Haenszel test. Data from 3 experiments with n=5/group. **d**. Volcano plot of differentially regulated genes in CD200 KO tumors vs. WT tumors. Red data points are genes with statistically significant (adj p value<0.05) and fold-change>2. n=3 mice. **e**. Network analysis of Gene Set Enrichment Analysis (GSEA) in CD200 KO tumor cells as compared with WT tumors. Network show pathways altered in CD200 KO tumors. Pathways positively enriched in CD200 KO tumors are in red. Pathways negatively enriched in CD200 tumors are in blue. Dot size represents enrichment score. Statistically significant pathway enrichments were corrected for false discovery rate (FDR) using Benjamini-Hochberg test. **f.** Top-15 most statistically positively (red) or negatively (blue) enriched pathways in CD200 KO tumors. **g**. Network analysis for Gene Ontology (GO) Biological Processes positively associated with CD200 expression in human glioma tumors (from TCGA). Statistically significant gene correlation and pathway enrichments were corrected for false discovery rate (FDR) using Benjamini-Hochberg test.

**Figure 3:**
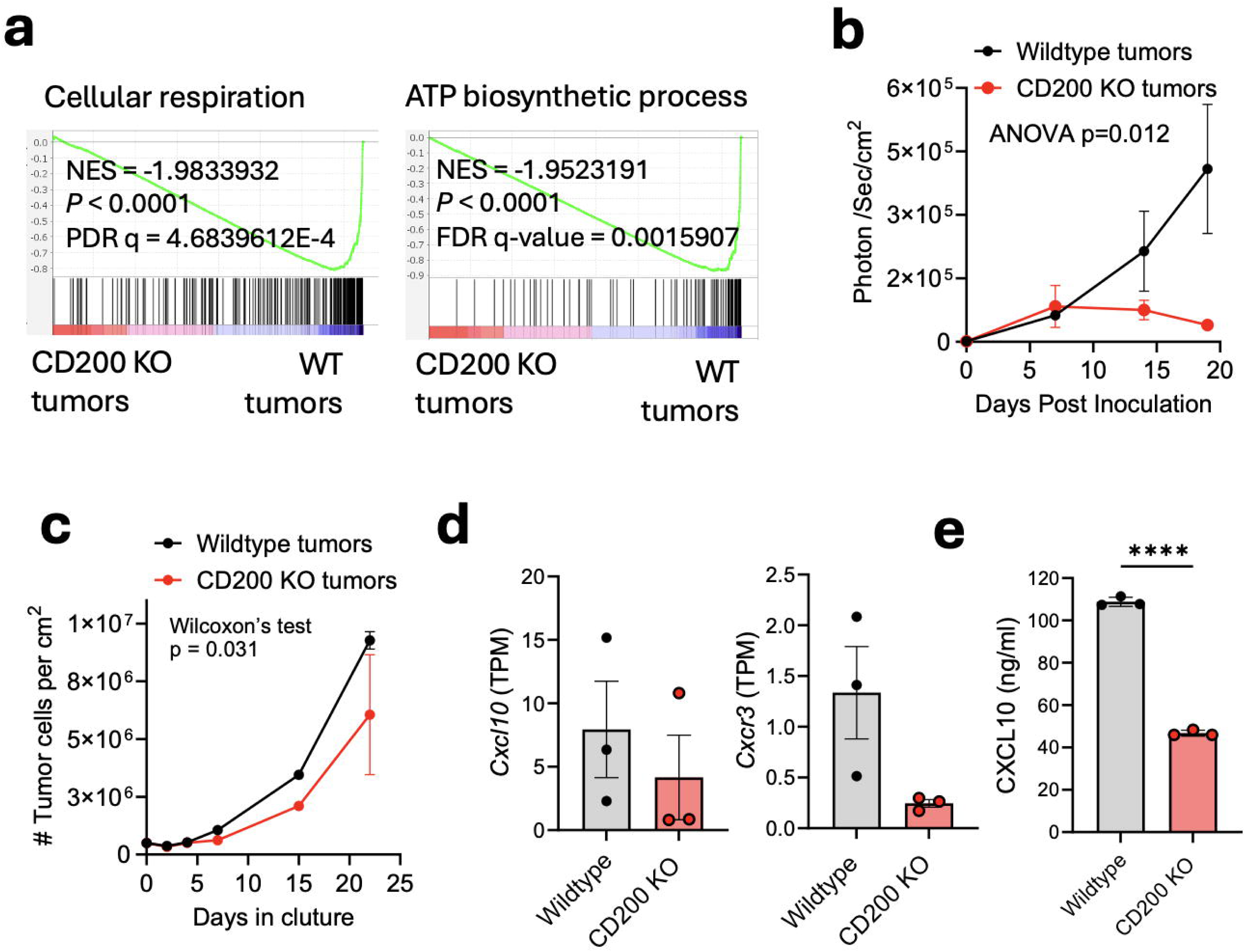
CD200 KO tumors demonstrate reduced cell numbers. **a**. Gene Set Enrichment Analysis (GSEA) of cellular respiration and ATP biosynthesis in CD200 KO tumors. **b**. Luciferin detection of WT and CD200 KO tumor cell in vivo. Data from 3 experiment. P value was determined using a one-way ANOVA test. **b**. Number of WT and CD200 KO glioma cells in culture. Data from 3 experiment. P value was determined using Wilcoxon test. **d**. Cxcl10 and Cxcr3 RNA expression (TPM) in wildtype and CD200 KO tumors. Data points show values of individual tumors. Student’s t test with Welch correction. **e**. CXCL10 concentration (ng/ml) in supernatant of WT and CD200 KO glioma cell cultures. Student’s t test with Welch correction. p****<0.0001.

### CD200 KO GL261 cells demonstrate reduced tumor cell numbers and decreased CXCL10 expression

Our findings demonstrate that CD200 KO gliomas were rejected in vivo (**Fig. 3c**), and that this was associated with changes in pathways associated with tissue development and morphogenesis, as well as with a decrease in cellular respiration and metabolism (**Fig. 3e-g**). For instance, cellular respiration and ATP biosynthesis pathways were significantly downregulated in CD200 KO tumors compared with wildtype (WT) tumors (**Fig. 3a**), thus suggesting that tumor cell growth was halted by lack of CD200 in these tumors (28). Correspondingly, we noted that as compared with WT glioma cells, CD200 KO glioma cells were spontaneously rejected over time despite the number of cells inoculated (**Fig. 3b**). Furthermore, although CD200 KO glioma cells could proliferate in culture, CD200 was required for optimal proliferation as observed by higher cell numbers of WT glioma cells in culture over time (**Fig. 3c**).

The chemokine CXCL10 and its receptor CXCR3 have been implicated in glioma cell proliferation (29). Accordingly, CD200 KO tumors were associated with a trend of decreased *Cxcl10* and *Cxcr3* gene expression (**Fig. 3d**). This was reflected in significantly lower CXCL10 levels in the supernatant of CD200 KO glioma cell cultures (**Fig. 3e**). Together, these findings suggest that CD200 contributes to glioma progression and may promote their growth, potentially through modulating factors like CXCL10/CXCR3 that can influence glioma proliferation.

### CD200 from glioma cells impedes anti-tumor immunity

While CD200 KO glioma cells showed a moderate decrease in growth in vitro (**Fig. 3c**), this effect could not fully account for the substantial decrease in CD200 KO tumor growth in vivo (**Fig 3b**), thus suggesting that, in addition to the associated TME metabolic changes (**Figs. 2e-f, and 3a**), additional mechanisms likely play a role in the rejection of CD200 KO tumors, including regulation of anti-tumor immune responses (30). Along these lines, we noted upregulation in immune response-related pathways (**Fig. 2e-f**), including the interactions between lymphoid and non-lymphoid immune cells, increases in cytokine expression and signaling, and antigen processing and presentation (**Fig. 5a**). Accordingly, our work and that of others, has established CD200 as an immune-checkpoint molecule whose expression hinders anti-tumor immunity primarily through NK cells and T-cells (9,31–35). However, the impact of CD200, specifically expressed in glioma cells, on the immune response against glioma has not been previously studied. Thus, we assessed if the expression of CD200 in glioma cells could regulate anti-glioma immunity.

Treatment of CD200 KO glioma-bearing animals with dexamethasone, a potent glucocorticoid that suppresses immune cells (36), negates the survival benefits associated with CD200 KO tumors (**Fig. 4b**). CD200 KO tumors expanded rapidly in vivo when animals were treated with dexamethasone (**Fig. 4c**). This suggests that the mechanism by which dexamethasone influences CD200 KO tumor cell numbers in vivo was not through directly altering tumor cell behavior (e.g., proliferation or growth), but more likely through suppressing the anti-tumor immune response.

**Figure 4:**
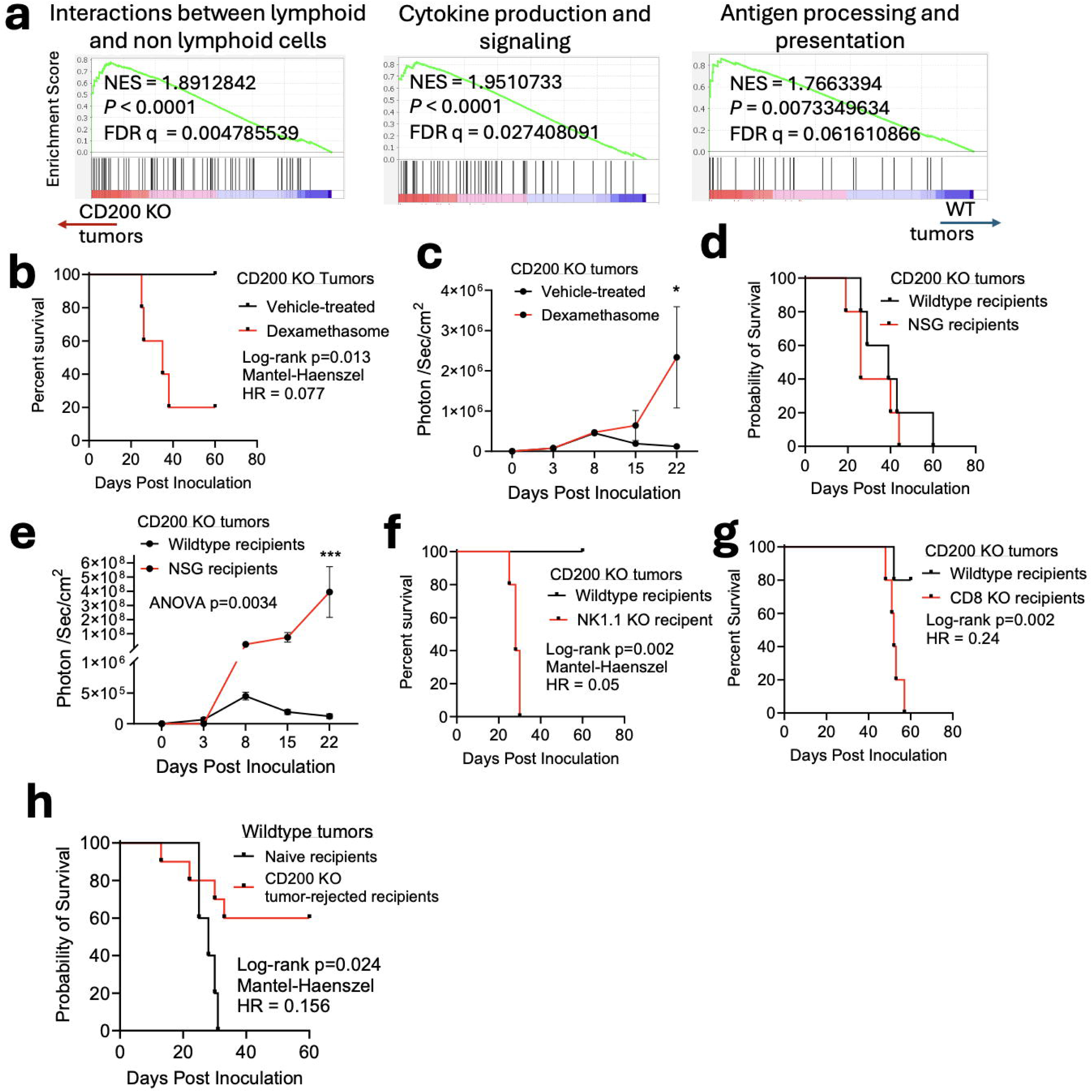
Spontaneous rejection of CD200 KO tumors is dependent on an intact immune system. **a**. Gene Set Enrichment Analysis (GSEA) of top 3 most enriched immune-related pathways in CD200 KO tumors. **b**. Kaplan–Meier survival curves of CD200 KO gliomas inoculated in mice following treatment with dexamethasone. p values were determined using a log-rank test. Hazard ratio was determined using Mantel-Haenszel test. Data from 3 experiments with n=5/group. **c**. Luciferin detection of CD200 KO tumors in mice treated with dexamethasone, as in b. Data from 2 experiment with n=5/group. Data points show mean with SEM. P value was determined using a one-way ANOVA test and post-hoc Tukey’s multiple comparisons test. * p<0.05. **d**. Kaplan–Meier survival of CD200 KO gliomas inoculated in immunocompromised (NSG) mice or immunocompetent mice (WT). P values were determined using a log-rank test. Hazard ratio was determined using Mantel-Haenszel test. Data from 3 experiments with n=5/group. **e**. Luciferin detection of CD200 KO tumor cells inoculated in immunocompromised (NSG) or WT recipients. Data from 2 experiment with n=5/group. Data points show mean with SEM. P value was determined using a one-way ANOVA test and post-hoc Tukey’s multiple comparisons test. *** p<0.001. **f**. Kaplan–Meier survival of CD200 KO gliomas inoculated in the NK deficient (NK1.1 KO) or WT recipients. P values were determined using a log-rank test. Hazard ratio was determined using Mantel-Haenszel test. Data from 3 experiments with n=5/group. **g**. Kaplan–Meier survival curves of CD200 KO gliomas inoculated in CD8 T-cell-deficient (CD8 KO) or WT mice. P values were determined using a log-rank test. Hazard ratio was determined using Mantel-Haenszel test. Data from 3 experiments with n=5/group. **h**. Kaplan–Meier survival curves of WT gliomas inoculated in naïve mice or mice previously inoculated with cD200 KO tumors following regression. P values were determined using a log-rank test. Hazard ratio was determined using Mantel-Haenszel test. Data from 2 experiments with n=5/group

**Figure 5:**
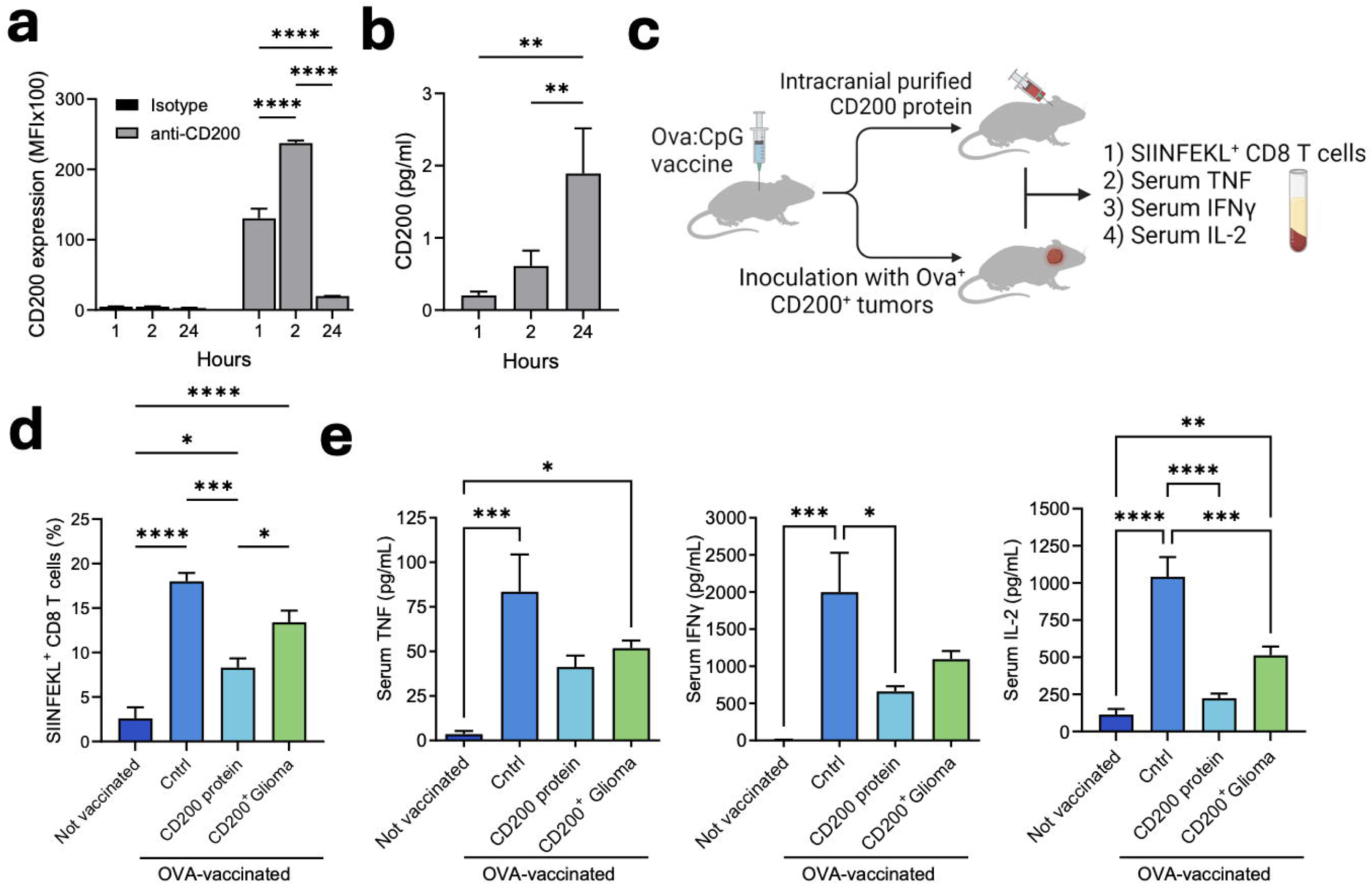
Glioma-derived CD200 impedes T-cell immunity. **a**. CD200 cell-surface protein expression on GL261 cells in culture over time. **b**. Concentration (pg/ml) of CD200 in supernatant of glioma cells in culture over time, as in e. P values were determined using one-way ANOVA and post-hoc Tukey’s multiple comparisons test. * p<0.05; ** p<0.01; *** p<0.001, **** p<0.0001. **c.** Schematic presentation of mice immunized with OVA protein emulsified in CpG followed by treatment with intracranial purified CD200 protein or inoculation of Ova-expression GL261 cells. **d**. Frequencies of OVA_257-264_ specific (SIINFEKL) T-cells from cervical lymph nodes from mice at day 21 post OVA vaccination. **e**. Serum concentration (pg/ml) of TNF, IFNγ, and IL-2 in mice at day 21 post OVA vaccination. Bar graphs show mean with SD. P values were determined using a one-way ANOVA test and post-hoc Tukey’s multiple comparisons test. * p<0.05; ** p<0.01; *** p<0.001, ****<0.0001. Data from 2-3 experiment with n=5/group.

To further investigate if an intact immune system was required for CD200 KO tumor clearance, we inoculated immunodeficient mice (NSG) or immunocompetent mice (WT) with CD200 KO glioma cells. In the immune-compromised setting, CD200 had no significant effect on survival (**Fig. 4d**). Furthermore, a significant increase in CD200 KO tumor cell numbers was seen when inoculated in NSG recipients compared to WT recipients (**Fig. 4e**).

Notably, while mice with a functional immune system spontaneously rejected CD200 KO tumors, those lacking CD8 T-cells (CD8 KO recipients) or with halted NK cells (recipients treated with anti-NK1.1 mAbs) rapidly succumbed following inoculation (**Fig. 4f, g**). Together these experiments indicate that the expression of CD200 by glioma cells could impede NK cells and CD8 T-cells anti-glioma activity which are pivotal for the rejection of CD200 KO tumors. Additionally, when CD200-expressing tumors were inoculated in mice that previously rejected CD200 KO tumors, we observed significant protection from a WT tumor recurrence (**Fig. 4h**). These data suggest the establishment of an immune memory in mice previously exposed to glioma lacking CD200. Taken together, these data support the critical role of CD200 in impeding anti-glioma immunity, including the establishment of T-cell memory.

### CD200 from glioma cells hinders circulating anti-tumor T-cells

Given the evidence that CD200 plays a role in tumor escape, we next wanted to assess the effect of secreted CD200 on antigen-specific T-cells and peripheral immunity. To determine if GL261 cells can secrete CD200, we cultured GL261 cells and determined the tumor cell surface-bound expression of CD200 as well as its levels in the supernatant over time. We observed that CD200 expression levels on gliomas were increased initially (1-2 hours), but significantly dropped after 24 hours in culture (**Fig. 5a**). This coincided with the significant increase of CD200 in the supernatant at this timepoint (**Fig. 5b**). Together, our data support that CD200 can be shed from mouse glioma cells.

We next vaccinated mice with ovalbumin following intracranial treatment with purified CD200 protein or with inoculation with CD200-expressing (i.e., WT) glioma cells (GL261) which also express the Ova protein (**Fig 5c**). These were compared to both unvaccinated animals and ova-vaccinated animals not receiving any treatment. Our study demonstrated that treatment with soluble CD200 protein or inoculation of mice with CD200-expressing gliomas, significantly reduced the frequencies of peripheral SIINFEKL-specific CD8 T-cells induced by vaccination (**Fig. 5d**). Additionally, treatment with soluble CD200 protein or inoculation of mice with CD200-expressing gliomas after vaccination with Ova, reduced serum levels of proinflammatory cytokines, including TNF, IFNγ, and IL-2 (**Fig. 5e**). Together these data suggest that glioma-specific expression of CD200 negated the priming and/or expansion of antigen-specific T-cells and may affect anti-glioma T-cell functions in dLNs, potentially through the shedding of CD200 protein into the circulation (10).

### CD200 protein increases in the serum of brain tumor patients undergoing tumor-cell lysate vaccine immunotherapy

Last, to evaluate if CD200 is present in the serum of brain tumor patients and potentially shed from tumor cells, we utilized blood samples collected from eight recurrent malignant brain tumor patients (six recurrent GBM, one ependymoma, and one medulloblastoma) who participated in a clinical trial assessing subcutaneous vaccination with dendritic cells loaded with allogeneic tumor lysate (GBM6-AD) (37). We conducted an analysis of CD200 protein levels in these serum samples and assessed their association with immune responses and T cells, an investigation not performed in the original published manuscript (37). Serum was collected in 4-week intervals up to 12 weeks, and clinical response was assessed by MRI (37) (**Fig. 6a**). As shown in **Table 1**, while the treatment demonstrated efficacy in prolonging progression-free survival (PFS), it did not confer a significant improvement in overall survival (OS), suggesting a transient therapeutic benefit. Half of patients (4/8) did not demonstrate a clinical response, and these patients also displayed no immune response and had elevated levels of regulatory T-cells (3/4). Three patients (3/8) had stable disease associated with immune response and low regulatory T-cells. One patient had a partial response associated with both a positive immune response and high regulatory cells (**Table 1**). Analysis of changes in serum levels from baseline (week 0; time of vaccine) across all patients showed that CD200 protein increased over time as tumor recurred (**Figs. 6b-d**). Together these finding demonstrate that CD200 can be found in the blood circulation of brain tumor patients, may increase overtime, and suggest that secreted CD200 levels might be associated with poor response to immunotherapy.

**Figure 6:**
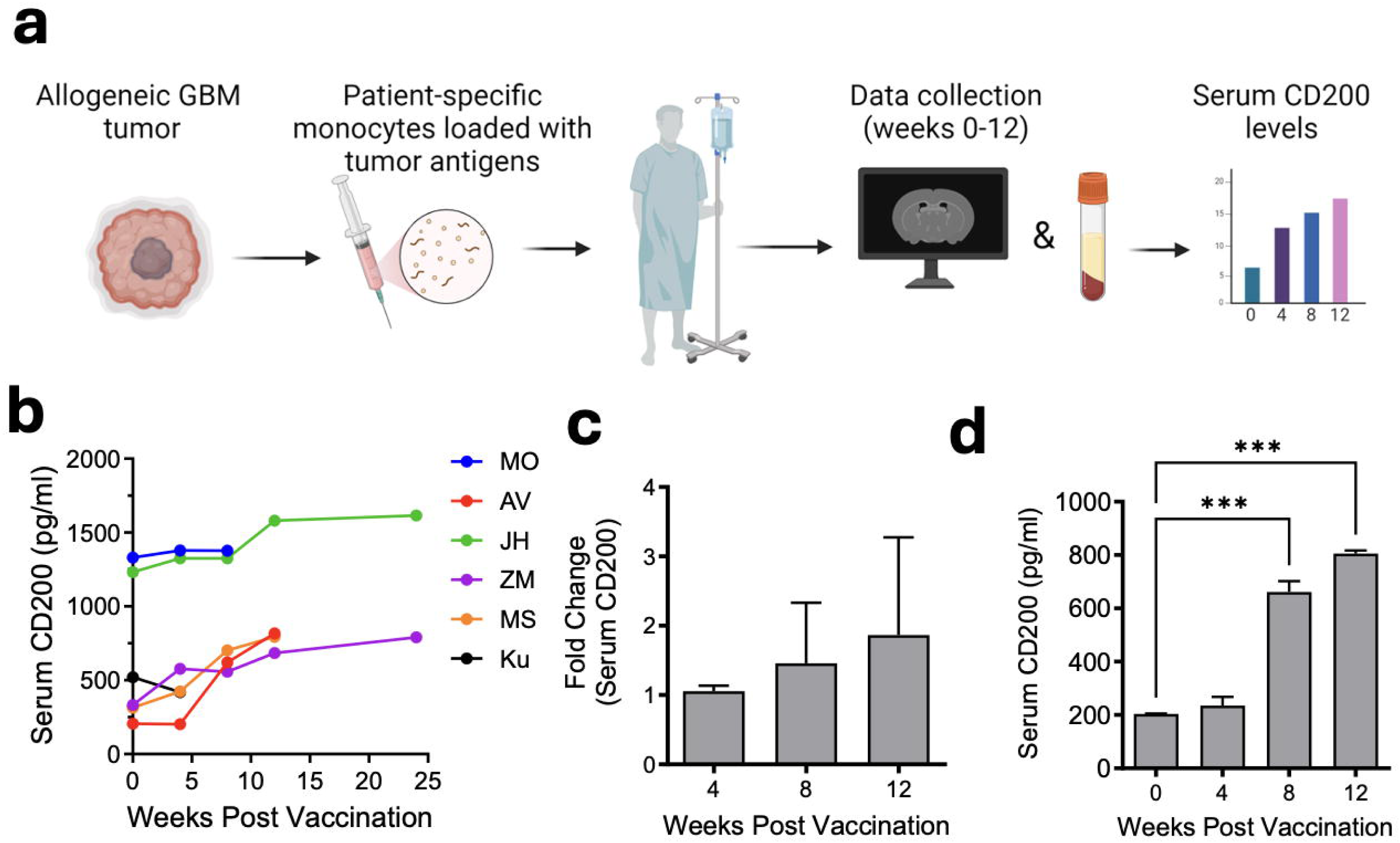
CD200 protein is shaded from tumor cells and is elevated in serum of glioma patients upon recurrence. **a**. Schematic representation of dendritic cell vaccine trial in patients. **b**. Serum concentration (pg/ml) over time of CD200 in patients following treatment. **c**. CD200 fold change in serum at weeks 4 to 12 compared to baseline. Data show mean with SEM. **d**. CD200 concentration (pg/ml) at weeks 0, 4, 8 and 12 post-vaccination in serum of a patient with stable disease and good outcome (patient ZM). Data show mean with SEM.

## Discussion

CD200 protein is primarily expressed within the central nervous system, however, CD200 is highly expressed in a variety of human tumors (9). The suppressive effects of CD200 can be mediated through interaction with the inhibitory receptor (CD200R1) on immune cells within the dLN or through interactions on the T-cells or tumor cells within the TME (17,38,39). While the exact mechanism remains unknown, we propose that CD200 works in concert with the dLN as well as on immune cells and tumors within the TEM. Previous studies and data reported here demonstrated that immune cells within the dLNs are susceptible to the suppressive effects of CD200 protein (9,31). Here we demonstrate that direct inoculation of CD200 protein into a normal brain demonstrates comparable suppressive responses to those seen in CD200^+^ glioma-bearing mice.

Within the glioma TME, several cell types can express CD200 and/or CD200R1 at various levels, including tumor cells, different myeloid cells (microglia/monocytes/MDCSs), lymphocytes (B cells and T-cells at various activation and differential states), and endothelial cells (9,11,40,41). However, it is unclear whether these cells are derived within the TME, originate within the dLNs, or differentiate upon interaction with the CD200 expressing tumor-associated vascular endothelial cells prior to entry into the TEM as we previously reported (31). Therefore, future work should focus on elucidating the effects of CD200 on tumor-related immune cells at different locations, both within and outside the TME.

Recently, mechanisms described for CD200 in tumorigenesis demonstrated that a 19 amino acid intracellular CD200 tail can be cleaved by γ-secretase resulting in the release of a CD200 tail fragment capable of nuclear translocation and affected cell proliferation (42,43). However, as tumor cells can also express the CD200R1, the role of the intracellular CD200 tail and CD200/CD200R1 on the TME and tumor cell functions should be characterized in future studies. Nevertheless, the role of CD200 acting directly on tumor cells may play a role in the induction of an immunosuppressive environment both in the TEM and within the dLN. Here we show CD200 KO tumors were associated with changes in immunological pathways as well as tumor cellular responses. CD200 could alter tumor function via downregulation of cell numbers and altered production of inflammatory mediators such as CXCL10, as we noted in vitro. CXCL10 can exhibit pleiotropic effects on a wide range of biological processes, including immunity, angiogenesis, and organ-specific metastasis of cancers, which is proposed to be a promising target for immunotherapy (44). Thus, we postulate that glioma-specific CD200 plays a pro-tumorigenic role via the engagement of CD200R1 in both tumor cells and immune cells, acting through direct tumor and immune-cell mechanisms.

Analysis of CD200 RNA expression within gliomas establishes that CD200 is expressed across all major histology groups and within different subgroups, which is in line with our prior report (9). Stratification of patients into risk groups based on the expression of CD200 revealed that low CD200 expression on tumors was associated with poor prognosis of patients, including worse overall survival rate and poor disease-free survival. Although it appears counterintuitive that patients with high tumor CD200 expression have better survival outcomes, this may be related to differences in CD200 shedding due to alterations in cleavage protein function and thereby lesser release of soluble CD200. In support of this view, prior reports have demonstrated that CD200 can be shed from tumor cells through cleavage by ADAM28 or metalloproteinases including MMP3 and MMP11, resulting in a biologically-active soluble CD200 ectodomain fragment (30,35). Furthermore, soluble CD200 can be found in serum of patients and may have immunoregulatory functions, such as NK cells and T-cells suppression (45–48). Hence, tumors can affect antitumor immune responses both through regulation on the enzymatic activity of proteases within TME, or through the increase levels of CD200 expression.

It has been largely accepted that CD200-CD200R1 promotes a pro-tumorigenic response, resulting in cancer progression; although, recent reports have also suggested an anti-tumorigenic role for CD200/CD200R1 (12,32,49,50), suggesting that CD200 may signal through activation receptors in addition to its inhibitory receptor CD200R1. Indeed, a few CD200 activation receptors (CD200AR) have been described to be predominantly expressed on immune cells of both myeloid and lymphoid origin (8). We posit that the anti-inflammatory effects of CD200 are due to proteolytic cleavage of the CD200 protein, exposing and/or releasing peptide sequences capable of binding the CD200ARs.

Here we demonstrate that mice bearing CD200 KO gliomas spontaneously rejected nearly all tumors. This rejection was immune-mediated, as mice lacking NK cells or CD8 T-cells CD200 KO tumor cells failed to control tumor growth. The observed suppression of T-cell and NK cell activity in glioma by CD200 in our study is consistent with findings in other cancers (13–16), further supporting its role in tumor immune evasion. Interestingly, our findings suggest that direct inoculation of CD200 into the CNS, even in the absence of a tumor, can cause immunosuppression of CD8 T-cells in LNs, mirroring the effects seen in tumor-bearing animals. This suggests a potential direct role for CD200 in immune regulation of T-cell priming or expansion, independent of tumor presence.

Taken together, our data suggest that glioma-derived secreted CD200 plays a key role in the pathogenesis of glioma through the regulation of anti-tumor immunity outside the TME, at least partially. We suggest that CD200 protein is capable of binding to uncharacterized activation receptors, and these have major implications for the development of the next-generation immune checkpoint blockade agents. Overall, our findings significantly advance our understanding of glioma-induced immune suppression via CD200, paving the way for the development of innovative immunotherapies.

## Acknowledgments

This work was supported by the Randy Shavers Research and Community Fund, Humor to Fight the Tumor, Children’s Cancer Research Foundation, and NCI Center for modeling tumor cell migration mechanics P01CA254849 and U54 CA210190 (MRO). This work was also supported by the UPMC Children’s Hospital of Pittsburgh Foundation (IR, GK), and the Walter L. Copeland funds of the Pittsburgh Foundation (IR). This research was supported in part by the University of Pittsburgh (Pitt) Center for Research Computing (CRC), RRID:SCR_022735, through the resources provided. Specifically, this work used the HTC cluster, which is supported by NIH award number S10OD028483. Schematic Figure 2a was generated using Biorender.com.

## Data Availability Statement

The sequencing data of GL261 CD200 KO and GL261 WT tumors generated in this study are available upon request from the corresponding authors. The analysis of adult glioma (LGG and GBM) RNAseq data in this study utilized, either partially or entirely, data produced by the TCGA Research Network (23,51) from https://www.cancer.gov/tcga, accessible on the GlioViz (http://gliovis.bioinfo.cnio.es/) (24) and GEPIA2 (http://gepia2.cancer-pku.cn/) (25) platforms .

## Author contributions

Conceptualization: M.G., C.L.M., M.G.C., G.K., M.R.O.; Data curation: I.R., A.A.M., R.E.S., B.M., C.T.S., A.M., A.D., E.A.M., M.N., F.P.M.; Formal Analysis: I.R., A.A.M., M.G.C., G.K., M.R.O.; Funding acquisition: W.B.E., M.G., C.L.M., M.G.C., G.K., M.R.O.; Investigation: I.R., A.A.M., E.A.M., F.P.M., M.G.C., G.K., M.R.O.; Project administration: R.S., C.T.S. M.R.O.; Resources: W.B.E., C.L.M., M.G.C., G.K., M.R.O.; Supervision: M.G.C., G.K., M.R.O.; Visualization: I.R.; Writing – original draft: I.R.; Writing – review & editing: I.R., M.N., C.L.M., S.C.F., M.G.C., W.B.E.,G.K., M.R.O.

